# Recent trends show a 20 year decline in global biological invasion rates

**DOI:** 10.64898/2026.11.13.766460

**Authors:** Brian Leung, Sami Sugiarta, Morgane Henry, Elizabeta Briski, Ross N. Cuthbert, Franz Essl, Núria Roura Pascual

## Abstract

Biological invasions are costly economically and environmentally. Despite international agreements to mitigate invasions, we have limited understanding of recent trends - the last two decades of invasion records have been typically excluded from analyses, given potential discovery and reporting lags. We use simulations to show discovery lags do not cause spurious recent declines, but reporting lags can do so. Using global empirical patterns, we estimated reporting lags caused up to a 94% underestimation of recent invasion rates. After controlling for these lags, our model estimated that the invasion rate accelerated at 1.5%/yr between 1970 and 2003 up to a maximum of ∼650 species/yr, but has since declined to ∼150 species/yr in 2020. Thus, for global trends, our analyses indicate that observed empirical declines are likely real and that lags cannot explain them, potentially suggesting that invasion policies have been effective.

## Introduction

Biological invasions are one of the primary drivers of biodiversity loss (Molot et al. 2017, Roy et al. 2024) and cause economic costs in the trillions of US Dollars (Diagne et al. 2021). These effects have been projected to increase, both due to changing environmental conditions (e.g., climate change, Omer et al. 2026), and increasing trade, which can introduce organisms to new regions (e.g., shipping: Sardain et al. 2019, Evans et al. 2025). Simultaneously, international agreements under the Kunming-Montreal Global Biodiversity Framework aim to reduce invasion rates by 50% by 2030, in order to safeguard biodiversity for future generations (CBD 2022). Beyond such aspirational future goals, numerous existing policies to prevent new invasions have already been enacted in recent decades. For instance, for marine invasions, the International Marine Organization’s Ballast Water Management Convention entered into force in 2017, but first started voluntary ballast exchanges in the late 1980s to early 1990s (IMO 2004). The International Sanitary and Phytosanitary Measures (ISPM15) requires the treatment of wood packing material associated with international trade – another primary pathway of entry of damaging insect pests (IPPC Secretariat 2009). At regional scales, the EU implemented a wide set of policies to mitigate the movement and impact of biological invaders (EU Regulation 1143/2014, European Commission 2014). While recent studies have shown that such policies reduce the rate of invasions (Canelles et al. 2025), often projections do not take recent policies into account (e.g., Sardain et al. 2019), or use historical invasion rates for calibration (e.g., Brockerhoff et al. 2014).

Given the relevance of species invasions, understanding changing patterns across the world is of fundamental and applied interest. Empirical data exist to potentially help, but these are typically compilations of invasion records across space and time, and are notoriously challenging to interpret (Brock & Daehler, 2025). Most prominently, lags have been identified as a source of bias, although previous work has primarily focused on discovery (detection) lags, defined as the time elapsed between the actual invasion and its discovery (Solow & Costello 2004, Robeck et al. 2024). Since discovery can occur years after invasion, the true invasion date is generally unknown (Fig 1, see Glossary in Supplementary Table S1). However, discovery does not necessarily mean that an invasion is immediately reported or that information about it becomes publicly accessible (Faccenda & Daehler, 2025, Macêdo et al., 2026). This introduces an additional delay, referred here as the reporting lag, defined as the time between discovery of an invasion and its public reporting. Moreover, only a fraction of reported invasions in the world will ultimately be included, constituting database completeness. Thus, it remains an open question as to whether we can use compilation databases to assess invasion trends, while recognizing that such databases are our primary means of doing so.

**Figure 1.**
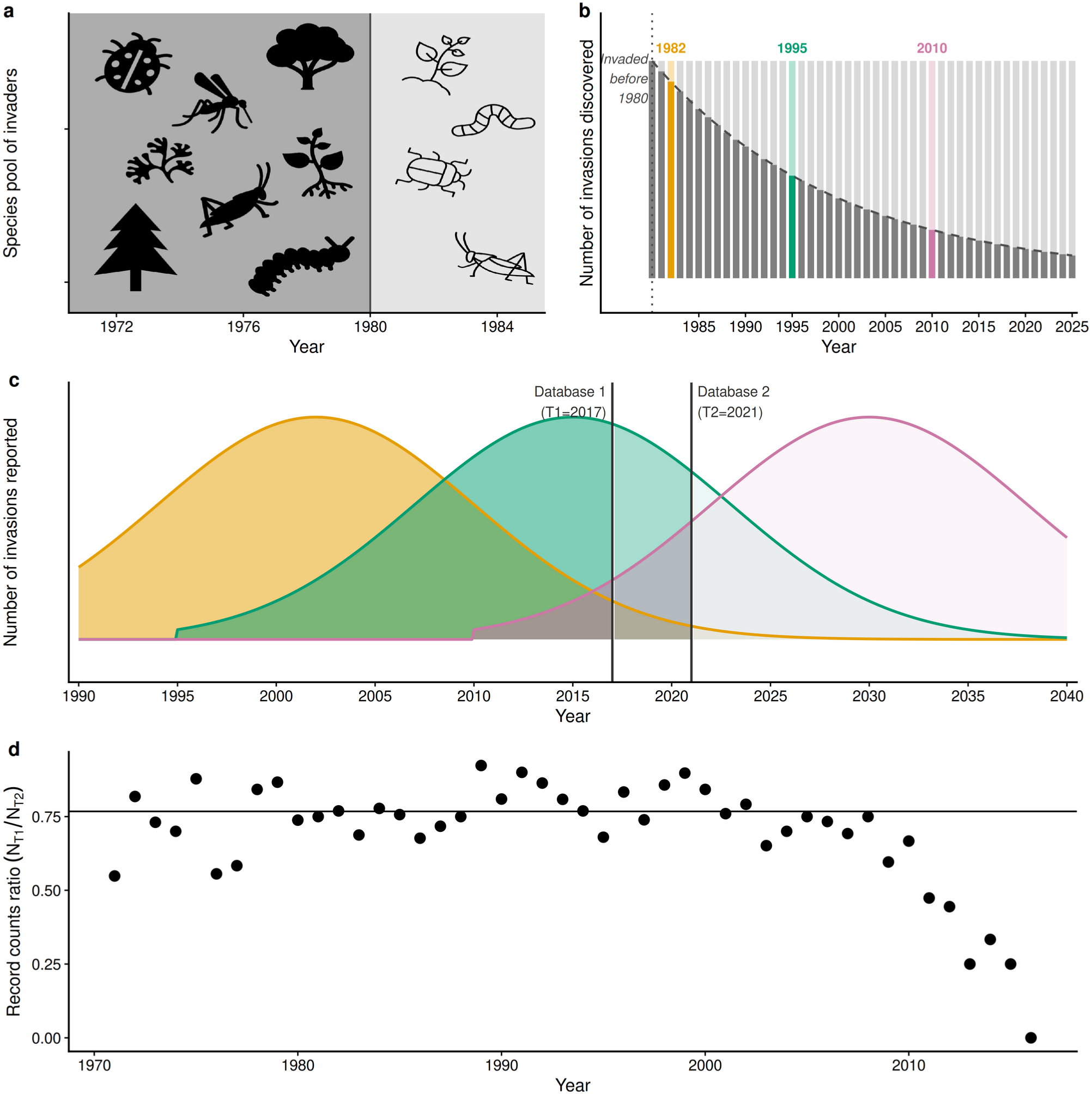
Process of First Records discovery and reporting in a database. *Panel a*: a pool of invasions that occurred across time. For illustration, invasions separated before and after 1980. *Panel b:* discovery occurs in years following invasion (i.e., discovery lag). Thus, discoveries in any year are comprised of invasions from the previously undiscovered pool. For simplicity, the pool of invasions and discovery rate are shown as constant, but more realistically, these can change over time. Dark bars - discoveries from pool of pre-1980 invasions; light bars - discoveries from post-1980 invasions. Discovered species removed from pool (dark bars decline over time). New invasions added to pool - discoveries increasingly reflect these new invasions (light bars increase over time). If more species are added to the pool than are discovered, the size of the pool grows, as does the subsequent rate of discoveries. As these balance, the rate of discovery approaches the rate of invasion. *Panel c: R*eporting occurs in years following discovery (i.e., reporting lag), resulting in distribution of reportings for each discovery date. Three discovery dates and distributions are shown (yellow – 1982, green- 1995, and pink – 2010; corresponding to labeled bars in *Panel b*). Only reports before a database release date can be included (i.e., cumulative area under curve to the left of vertical line, e.g., for Database 1). Subsequent updated versions of the database (vertical line for Database 2) can include additional invasions discovered *prior* to release of Database 1, since more time has elapsed for reporting. *Panel d*: Taking the ratio of records between databases (N_Database1_/N_Database2_) for each year should theoretically yield a curve where ratios are most different in years closer to the release date of Database 1, and becomes more equal for older discoveries, as cumulative distribution increasingly occur in Database 1. Additionally, databases are invariably incomplete (i.e., not all reported species in the literature occur in any given database) and additional sources can be included in subsequent databases (increasing completeness), resulting in an asymptote (horizontal line)which estimates relative completeness. This will be the logical basis of the “*reporting ratio”* model developed in this article.

Importantly, lags for discovery, reporting and database completeness can have distinct effects, and thus are necessary to separate (see Glossary, Supplementary Table S1). For instance, database completeness can reflect inclusion of whole new sources of data (Seebens et al. 2018), and thus updated versions of a database can increase the number of records across all time points. In comparison, for reporting lags, we should see temporal patterns, with fewer records closer to the publication date of a given database (Fig 1). Further, for a given year more time will have elapsed for more recent databases, which in theory should result in different probabilities of being reported. This discrepancy between databases might allow estimation of the reporting lag. Moreover, while database completeness is driven by its curators, discovery and reporting reflect efforts within the larger invasion biology community; effects of changes in effort should also be considered. Regardless, we argue that elucidation of the effects of discovery and reporting lags, changes in database completeness and effort are necessary to understand trends in invasions and how these may be changing.

As a partial solution to these systematic biases, previous analyses have typically excluded the last couple of decades of invasions to account for lags, analyzing reports up to the early 2000s, which saw a continued increasing rate of invasions (e.g., IPBES, 2023; Seebens et al. 2025). However, we also expect to see the effects of alien species policies in this same time period, since most policies have been implemented in the last two and a half decades (IPBES, 2023). We argue that instead of censoring empirical patterns, it would be ideal to account for potential effects of temporal biases, to allow a fuller understanding of the available data and potential effects of policy interventions on global invasion trends.

Here, we provide an estimate of the most recent, critically important, invasion patterns using the First Records Database (Seebens et al. 2017) - the most comprehensive source on the first recording of alien species in different regions (i.e., countries, islands, federal states). To do so, we first examine theoretically the effects of discovery and reporting lags, database completeness and effort on apparent invasion trends. We then develop an approach to estimate invasion rates, using two versions of the database taken at different time points to separate competing phenomena of invasion rates, discovery and reporting lags, and completeness; and we show where they are theoretically able to capture underlying processes, and where they may fail to do so. Finally, we examine how regional and taxonomic biases may influence global estimates and the resulting invasion patterns.

## Methods

### Model formulation

The main components of the model are invasion rates (with a potential change point), and probabilities of discovery and reporting and database completeness in the First Records database, with model specification defined in equations 1-7 (also see Glossary in Supplementary Table S1).

We use an exponential model for invasion rate and modeled a potential change point at some time t_c_ (equations 1 & 2).

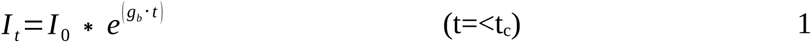

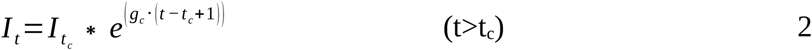

where I is the invasion rate (i.e., the number of invaders at a given time interval t), I_0_ is the intercept (initial invasion rate), g is the rate of change before (g_b_) and after (g_c_) the change point year (t_c_) (i.e., when t>t_c_), and I_tc_ is the invasion rate at the change point year. This model constrained the number of invasions so that it could not become negative.

After each invasion event, the species could be discovered over time (i.e., a discovery lag). We considered the possibility that probabilities of discovery during a given time interval could increase as an invasion progressed. In notation:

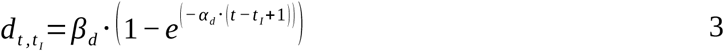

where d is the probability of discovery during time interval t, t_I_ is the time interval that a species invaded, and α_d_ and β_d_ are detection coefficients, to allow flexibility in how discovery probabilities change over time. From equation 3, it can be seen that the coefficient β_d_ determines the maximum probability per time interval (0< β_d_ ≤ 1), and α_d_ allows discovery probability to increase with time since invasion (α_d_ > 0) – higher values of α_d_ cause discovery probabilities to increase faster over time. At large α_d_, the probability of discovery asymptotically becomes a constant value β_d_ per time interval. As discovery probability decreases, discovery lag increases.

Thus, the expected number of discoveries at a given time interval t (D_t_), is determined by the number of invasions at each previous time interval, the probability that it was discovered during time t, and had not been previously discovered between the invasion date (t_I_) and t. In notation:

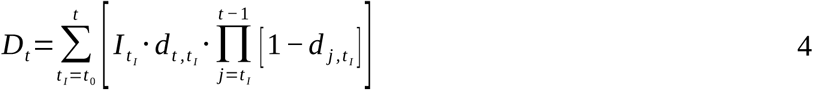

More simply, the discoveries at t depend on the probabilities of discovery and pool of previously undiscovered invasions (Figure 1b).

Similarly, a species could be discovered in a given year t_d_, yet remain unreported for several years (i.e., reporting lag). Here, we used a similar formulation and logic as for discovery (equation 3) to model reporting lags:

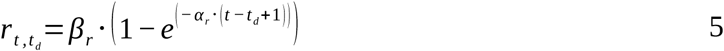

where *r_t , t_* is the probability of being reported in year t, if the species were first discovered in year t_d_, and α_r_ and β_r_ are reporting coefficients. Invasions must be discovered and reported, before they can possibly be included in a database.

In notation,

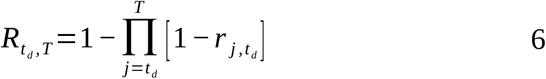

where *R_t , T_* is the cumulative probability of a species detected in year t_d_ , and also reported in the literature before the database released at time T.

Only a fraction of invasions that have been both discovered and reported will be included in a given database to form a “first record”. As such, we also include a coefficient (M_T_) to describe completeness of a database released at time T. Thus, we use M_T_ and equations 4 and 6 to calculate the expected number of records (*E_t , T_* ) of invasions discovered at time t_d_, reported and included in a database released at time T, according to:

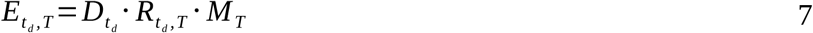

### Analytic approaches to fit data

The model above describes the process of invasion, discovery, reporting, and database entry (completeness) into the First Records database. The question is whether there is sufficient information in the observed counts to fit the underlying processes. To do so, we developed two complementary statistical models: “*reporting ratio model*”, to estimate reporting lags, and the “*record count model*”, to estimate invasion rate, after controlling for reporting lag. These are described below. We fitted model parameters using Maximum Likelihood, using R version 4.3.1 (R Core Team, 2023)

For the *reporting ratio model,* we propose that comparison of two versions of the database taken at different time points could provide the information needed to separate the competing phenomena of invasion rates. Specifically, by fitting the ratio of the number of reported species at each time interval, from two subsequent database versions (N_older_/N_newer_), we can control for the number of discoveries in each given year, because this quantity is shared by both database versions and therefore mathematically cancels out. Notably compilation databases are comprised of heterogeneous sources of records, and any temporal analyses using these (e.g., IPBES, 2023) implicitly assumes that aggregation across many sources “average out” this heterogeneity, to allow extraction of general trends. Likewise for comparisons of database versions. Adding sources to a database -i.e. improvement in database completeness - should result in a proportional difference in the number of records if the underlying trends are comparable. However, in contrast to general database completeness, reporting lags will theoretically cause a different (temporal) pattern - because the cumulative probability of reporting increases over time, and the newer database has had more time for reporting, the ratio between the number of records in the older and newer databases (N_older_/N_newer_) should increase over time, approaching a constant asymptote (Fig 1d). Thus, temporal changes in the ratio allow estimation of the reporting lag, whereas the asymptote allows estimation of completeness of the older databases relative to the newer one.

To construct the *reporting ratio model*, we used the following procedures:

The true numbers of invasion I_t_ and numbers detected for the first time at time t ( *D_t_* ) were unknown; only the number of records *O_t , T_* for each discovery date (t_d_) was known. However, the ratio of records (equation 7) from two databases taken at different time points (T1 and T2) would factor out the unknowns *D_t_* (and I_t_) and provide an estimate of temporal differences in reporting rates. Database versions also differed in their completeness (we assumed delistings did not dominate database differences). While we could not estimate the absolute values of completeness (M_T_ in equation 7), we could estimate the relative completeness of databases at T1 compared to the database at T2 (M_R_ = M_T1_ / M_T2_; 0 < M_R_ < 1), which was sufficient to account for differences in completeness. In notation:

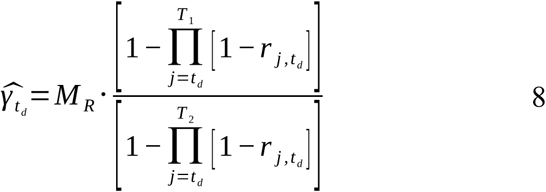

where *^γ_td_* is the ratio – the expected fraction of records in the database at time T1 compared to T2. To fit the data, we used *^γ_td_* as the probability in a Binomial likelihood, and compared differences in the observed record counts *O_t , T_* in T1 versus T2. In notation:

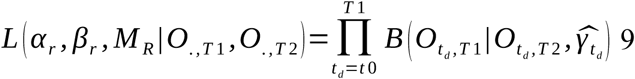

where B denotes the Binomial likelihood. The dot subscript . denotes all time points under consideration. O_td,T1_ acts as the number of “successes”, and O_td,T2_ acts as the number of “trials” in a Binomial distribution, at each time interval j, given the expected probability *^γ_td_* for each discovery date (t_d_). We used this likelihood (Equation 9) to fit parameters described previously: α_r_ and β_r_ (equation 5) to estimate reporting lags, and also M_R_ (equation 8) to account for differences in database completeness.

Next, to estimate the remaining parameters, we constructed our *record count model*. We fitted the combined model (equations 1-7), comparing expected *E_t , T_* _2_ versus observed *O_t , T_* _2_ numbers of records at each time interval. While we could fit observations from either database, we used the more recent database at T2, because it was more complete. We used the maximum likelihood probabilities *R_t , T_* _2_ for each time interval (equations 6, 8, 9). This helped with issues of identifiability, rather than fitting all parameters simultaneously, and allowed separation of reporting lags (probabilities) from change-points of invasion rate (equations 1 & 2). For this analysis, we could disregard database completeness (set M_T2_ = 1), since we were only modeling relative temporal trends *within* a database. We used a Negative-Binomial (NB) likelihood, given that we had discrete count data, and overdispersion in the First Records database empirical patterns. In notation:

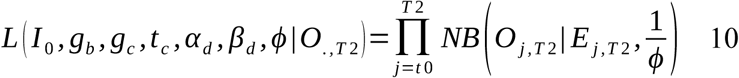

We used the version of NB parameterized using the expected number of records (E_j,T2_, equation 7) and an overdispersion parameter ϕ (using the alternative parameterization using “mu” in the dnbinom function in R). Additionally, we used an inverse dispersion parameter, such that the likelihood approached a Poisson distribution as ϕ goes to zero, and dispersion became infinitely large as ϕ approached zero.

### Simulation analysis of model accuracy

To evaluate the performance of our model, we used simulation analyses. We generated virtual datasets with known parameter values to examine the ability of the *reporting ratio model* and *record count model* described above to correctly estimate the parameter values. These analyses allowed us to assess model reliability across a broad range of invasion, discovery, and reporting scenarios (by varying parameter values).

In particular, we modeled 70 years, with a 20 year “burn-in” to allow equilibration from starting conditions. We simulated discovery and reporting parameters between: 0.05 ≤ α. ≤ 1.6 and 0.3 ≤ β. ≤ 1, which yielded initial probabilities ranging from 0.015 to 0.8. For reporting probability, we also simulated the relative completeness coefficient between 0.3 ≤ M_R_ ≤ 1. Initial generating invasion rates were increasing, simulated according to g_b_ ∼ N(μ = 0.05, σ = 0.03), with a minimum of g_b_ truncated at 0.001. To put concrete values to the parameters, at a value of g_b_ = 0.05, the number of invasions would double after ∼14 years. For the rate of change (g_c_) of invasions after a change point t_c_, values were simulated according to: g_c_ ∼ N(μ = 0, σ = 0.07), such that we modeled both increasing and decreasing rates after t_c_ . The timing of change-points was randomly chosen between 45 ≤ t_c_ ≤ 55 years (to match the time frame of the observed empirical declines in our analyses of the First Records database). For sampling distributions, we modeled a Negative Binomial distribution, with an overdispersion parameter 0.01 ≤ ϕ ≤ 10.24. In combination, this generated a wide range of potential relationships to examine the theoretical accuracy of our models, evaluated by how well our fitted parameter values correlated with the true parameters across all simulations.

### Theoretical effects of changes in effort

We examine the patterns generated by doubling and halving probabilities of discovery and reporting from a baseline (7%), for several scenarios: a short-term “pulse” of effort (e.g., due to a grant or short-term program) and a long-term “press”, where effort remains elevated or reduced. Additionally, we examine the consequences if effort changes between database versions. These are arguably the most relevant scenarios, given these may affect the estimates of the model, and our analyses provide diagnostics to identify when caution is warranted. An alternative approach could be to weight probabilities using secondary data sources (e.g., the GBIF database, GBIF.org 2025), which assumes that effort for invasion monitoring changes similarly to the secondary sources. However, we chose not to take this approach given uncertainty about this assumption (for instance numbers of new native records likely occur in very different conditions than new invasive species observations), and because correcting for the temporal increase in effort in GBIF (for instance), would only exacerbate the estimated decline. Thus, our approach is conservative.

### Analysis of First Records Database

To apply our modeling framework to empirical data, we used the “Alien Species First Records” database, first published by Seebens et al. (2017), which provides temporal information on first records of alien species. This database has been used to estimate invasion trends (e.g., IPBES, 2023). Further, there were multiple versions of the First Records database (2017, 2021, 2023, and 2025) which offered an opportunity to fit our *reporting ratio model* and to extract important information on reporting lags. Further, as recent invasions and their changes were our primary interest, we focus on invasions after 1970, to reflect contemporary patterns, and to balance comparability of reporting and discovery lags across the time series with having sufficient records of invasions. This date range provided sufficient data before and after apparent switch from increasing to decreasing numbers of new invasions reported, allowing statistical tests for a change point, after considering discovery and reporting lags.

Since the analysis compared multiple iterations of the database, we preprocessed the data to ensure comparability. Firstly, we used the countrycode R package to obtain the region and sub-region of the given locations, as defined by the UN (Arel-Bundock et al. 2018). Where the location could not be resolved, the data were entered manually. Secondly, we used the rgbif R package to resolve taxon names using the GBIF backbone taxonomy (Chamberlain et al. 2017). We discarded any taxa that could not be resolved down to the species level or whose match type was not one of “FUZZY” or “EXACT”. We then removed entries whose year was not in the range of consideration.

Beyond estimation of rates, we examined the ΔAIC. For the *reporting ratio model*, we tested for evidence of a temporal reporting lag against a null model which only included differences in completeness between databases. For the *record count model*, we tested for evidence of a change-point in invasion rate (in particular changing from increasing to decreasing rates), against a null model of a constant invasion rate. Finally, we tested whether inclusion of the reporting probabilities from the *reporting ratio model* improved the fit to the record counts, compared to model assuming no reporting lag. While lack of improvement does not necessarily imply absence of a reporting lag, since the change-point in trend could capture the empirical decline irrespective of the mechanism, an improved fit would provide additional support that the estimated reporting probabilities are meaningful. Conversely, lack of improvement would suggest caution in interpreting trends.

We first conducted the analysis using the entire dataset to identify aggregated patterns at a global scale. Subsequently, we repeated the analyses after separating the data by region and taxonomic group. This approach allowed us to assess whether the overall patterns identified at the global scale were consistent across different geographic regions and taxa, as well as to detect potential emerging or divergent dynamics that might otherwise remain masked in the aggregated analyses.

## Results

### Theoretical behaviour

We found that discovery and reporting lags had radically different consequences for invasion patterns. Based on our theoretical analyses, discovery lags do not cause apparent (false) declines, or changes from increasing to decreasing invasion rates (Fig 2a-d). Even when species are discovered with only a low probability, and recent invasions are likely to have been missed, the number of new discoveries does not decline.

**Figure 2.**
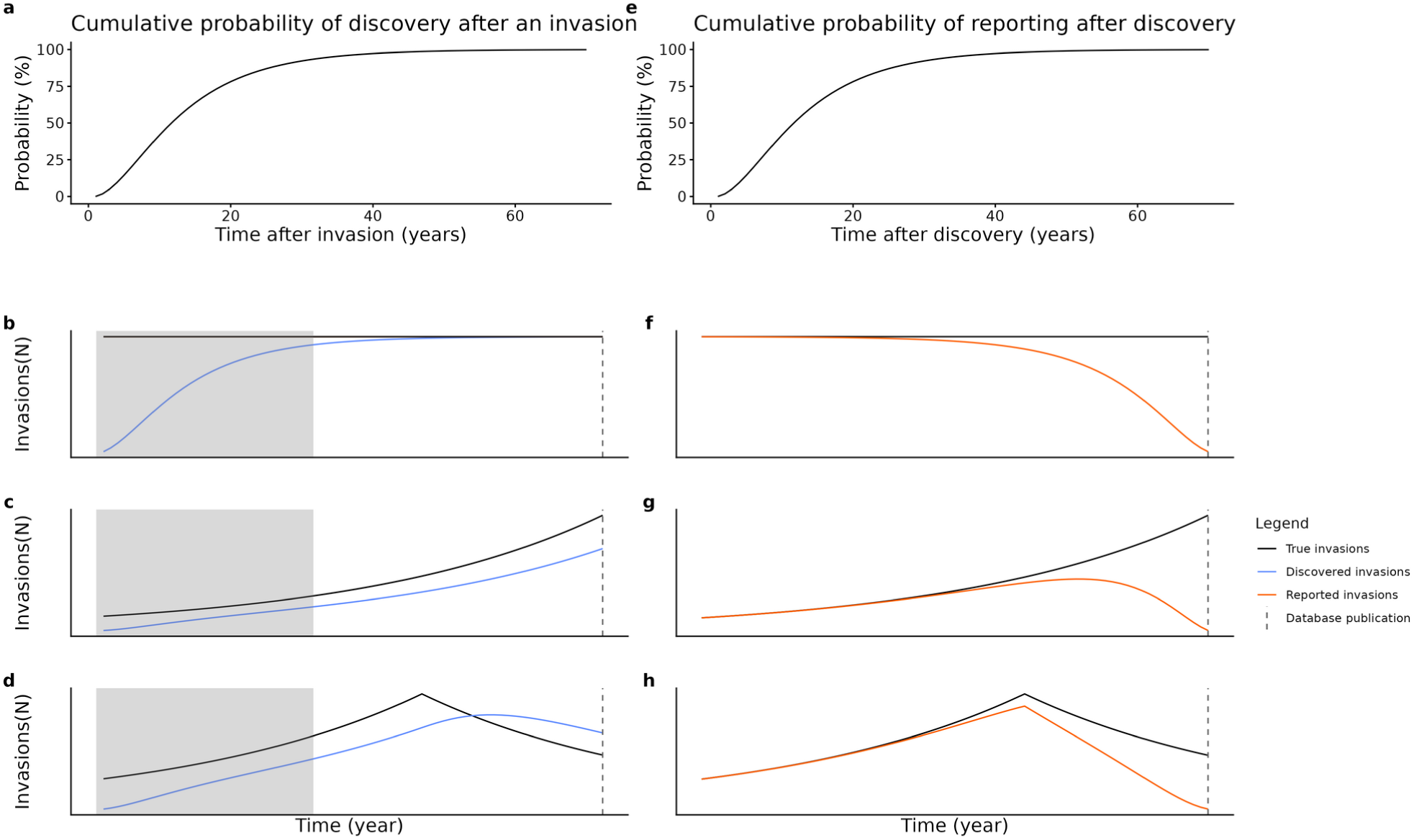
General effect of reporting and discovery lags. The top panels show the probability of discovery as a function of time since invasion (a), and probability of reporting as a function of time since discovery (e), as an outcome of the processes described in Figure 1 (conceptual diagram) and equations 1-7. Here we examine the effect of each type of lag separately, to highlight the differences in their relationships with underlying invasion rates (black lines), for constant (b, f) and increasing (c, g) invasion rates. Additionally, change points with increasing to decreasing invasion rates were shown (i.e., d, h). With discovery lags (panels b-d), there is an initial artifactual increase as the discovered numbers of invasions equilibrates (grey areas). After equilibration, the discoveries match the underlying invasion rate, although when rates change (d), discoveries show a lagged, smoothed transition. In contrast, with reporting lags (panels e-h), recent years will show artifactual declines, even for constant (f) or increasing (g) invasion rates, and therefore can be conflated with true declines (h).

This occurs because the number of discoveries in a given year will have contributions from invasions across prior years, from the pool of previously undiscovered species (Fig 1a,b). As each species is discovered, it is removed from the undiscovered pool. If the new invasions entering the pool exceed the numbers discovered, the pool grows and increases the subsequent numbers of discoveries. As the numbers of discoveries and new invasions reach an equilibrium, the rate of discovery approximates the rate of invasion. Note that equilibrium is reached even for low probabilities of discovery (but takes longer). Additionally, we note that there can be an artifactual *initial* increase in discoveries, while the pool size of alien species increases, until the invasion and discovery rates equilibrate (Fig 2b, grey area). Arguably, this will likely not be an issue for recent invasions, as invasions have been happening for many decades and centuries before, allowing time for equilibration. Instead, limitations caused by the discovery lag are two-fold: *(i)* the true absolute number of invasions in each year remains unknown; and *(ii)* the timing of estimated changes in invasion rate (i.e., a change-point) is biased towards later dates.

In contrast to discovery lags, reporting lags can result in apparent declines closer to the release date of a database, such that constant or even increasing invasion rates could still show a declining number of newly reported species invasions. As such, a correction for reporting lags is necessary to differentiate between reporting lags and truly declining invasion rates: it would be difficult to differentiate between reporting lags and truly declining invasion rates (Fig 2e-h), without a correction for reporting lags.

The difference in behaviour arises because of the information retained for discovery versus reporting lags. For discovery lags, each discovery year is comprised of species with unknown invasion year, reaching an equilibrium with the invasion rate as described above (Fig 1b). In contrast, for reporting lags, each discovery year has been recorded and can be identified separately; whether it is reported or not in a database is therefore simply the cumulative probability, as a function of the time between the date of discovery and the database release (Fig 1c).

### Theoretical analysis of parameter estimation

We first present simulation analyses of the *reporting ratio model*. If a reporting lag is detectable, we expect that there will be a temporal signal, with smaller ratio more recently (Nolder/Nnewer), increasing to an asymptote, which reflects differences in completeness (Fig 1d). We found that the approach theoretically allowed statistical estimation of both the probabilities of reporting (Supplementary Fig S1a,b), and relative differences in database completeness (Supplementary Fig S1c), used to simulate the data.

In contrast to reporting lags, there was no tractable approach to mathematically isolate discovery probabilities. While a discovery sub-model could be fit simultaneously with invasion rates, we found that we could not reliably estimate discovery probabilities (Supplementary Fig S2). However, provided reporting lags are accounted for, we show that we can set discovery lag to zero, and still obtain good estimates of the other parameters underlying the *record count model*, including the true underlying invasion rates (Supplementary Fig S1d,e) and the error distribution (Supplementary Fig S1f). This was because discovery rates asymptotically estimate the true invasion rates over time (as shown in Fig 2b-d). Although we could detect that a change in invasion rates had occurred (Supplementary Fig S1d,e), we note that the exact time estimate of the change-point date was unreliable (Supplementary Fig S1g). Thus, we exclude discovery lags from the remainder of analyses.

Practically, this simulation illustrates that robust analysis of recent invasion rates requires reporting lags to be accounted for, whereas discovery lags are relatively unimportant.

### Theoretical effects of changes in effort

When effort increased substantially causing discovery probability to increase, we observed a theoretical spike in new discoveries (Supplementary Fig S3), which might thus be used as a characteristic indicator of changed effort. If effort increased as a single, one-time event, the discovery rate would return to previous levels in subsequent time intervals (Supplementary Fig S3a). If this increased effort continued across time, the discovery rate would remain elevated for a time (potentially decades), but would gradually decline back to the true invasion rate (Supplementary Fig S3b), despite persistently higher effort. This is because higher rates of discovery deplete the pool of undiscovered species faster than new invasions are added, causing the rate of discovery to decline until equilibrium between invasion and discovery rate is reached. Conversely, if effort is reduced, the reverse pattern occurs, with the lower discovery probability causing a longer time to reach equilibrium again. Thus, sudden spikes in discoveries are more likely to reflect changes in effort than changes in true invasion rates; actual spikes in invasions tend to be smoothed out due to the averaging effect of discovery lags. Concomitantly, the occurrence of discovery spikes could be used as an indicator to separate changes in effort versus changes in invasion rates.

In comparison, for reporting lags, changes in effort do not have an obvious indicator within a single time-series, as these are determined by the cumulative probability of reporting between discovery and the database release (i.e., the exact timing of effort is less important). However, change between databases remains an important consideration, because in our approach reporting lag is calculated using the ratio of records between databases. If reporting probability increases in the latter database, the difference in number of reports also increases. Counterintuitively, this reduces the estimated reporting probability, inflating the reporting lag, and causing the invasion rate to be overestimated. Conversely, if effort is reduced, the difference between databases will decrease and the lag will appear shorter. In the most extreme case, the temporal signal of reporting lags can disappear entirely, such that reporting is estimated to be immediate. Consequently, the observed decline in alien species numbers over time may be erroneously attributed to true declines.

### Global patterns of biological invasions

We first compared the empirical patterns from the four versions of First Records databases (Fig 3). Examining the raw annual first records showed higher completeness for each subsequent version of the database, as expected (Fig 3a). Further examination of normalized annual invasion numbers (by dividing by the maximum within each database), showed several important characteristics. First, despite the heterogeneous sources added to each database, the increases across databases were proportional (Fig 3b) and relative changes across years were consistent after reporting lags. This finding is consistent with the assumptions of the model and represents an important prerequisite for interpreting global invasion trends. Second, there was no evidence that the empirical decline was due to reporting lags, which should result in declines shifting to later years with subsequent databases. Instead, the decline remained approximately constant beginning in year ∼2000, despite almost a decade between the earliest and latest databases. We note, however, that the recent 2023 and 2025 databases showed few records after 2020, with zero reported records after 2021 and 2022, respectively.

**Figure 3.**
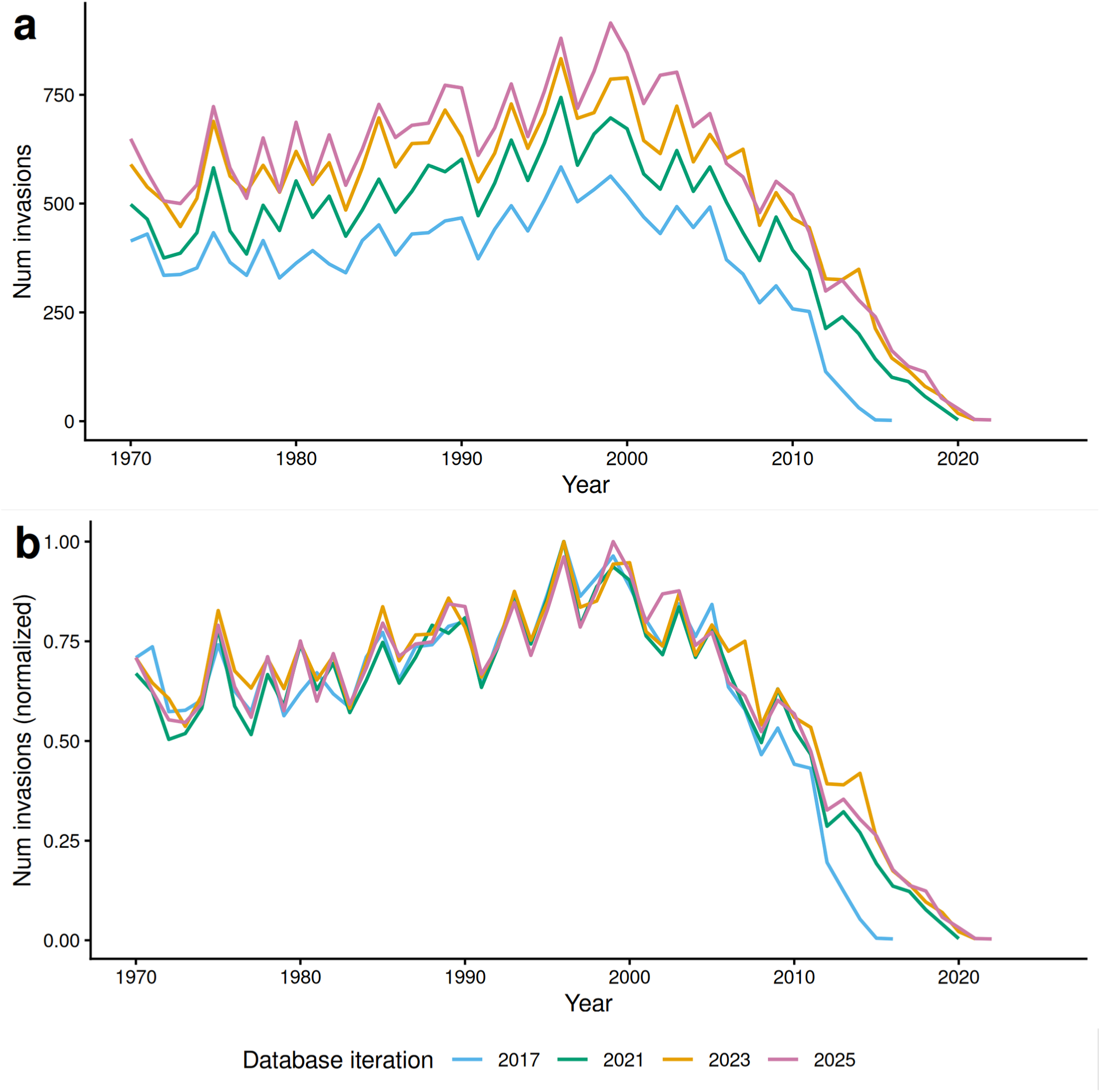
Annual empirical record count, for four versions of First Records database. a) Raw counts of number of invasion records over time. Higher counts in more recent databases reflect added sources and increased completeness. b) Records normalized by the maximum observed in that database version (between 1970 and 2025). Strong overlap between databases across earlier years shows that trends changed approximately proportionally between databases, despite considerable absolute differences between the earliest (2017) and latest (2025) databases. Count differences between databases in recent years show reporting lags.

More broadly, the 2023 and 2025 databases showed potential evidence that reporting probabilities were lower than the 2017 and 2021 versions. As discussed above, reporting should be disproportionately higher for a given year (approaching an asymptote over time), because later databases have had more time for reporting. This reporting-driven deviation is apparent for 2017 versus 2021, but 2023 and 2025 are both similar to 2021 (Fig 3b), suggesting relatively reduced reporting probability. Given that our theoretical analyses show that reduced reporting probability between databases results in overestimated rate of decline, we focus the remainder of our analyses on the 2017 versus 2021 databases.

Using our *reporting ratio model*, we found a clear temporal signal of reporting lags globally (Fig 4a). Our analysis suggests a lag of nine years since discovery for 95% of invasions to be reported. Additionally, there was a general difference in completeness between databases, with ∼20% fewer invasions recorded per year in general in the earlier database. For comparison, the ratio model estimated 92% fewer invasions in 2016 in the early database versus the latter database. This is broadly consistent with the empirical observations - for discoveries in 2016, two invasions were recorded in the 2017 database compared to 101 records in the 2021 database. Thus, the temporal effect of reporting lag was apparent far beyond the general difference in database completeness.

**Figure 4.**
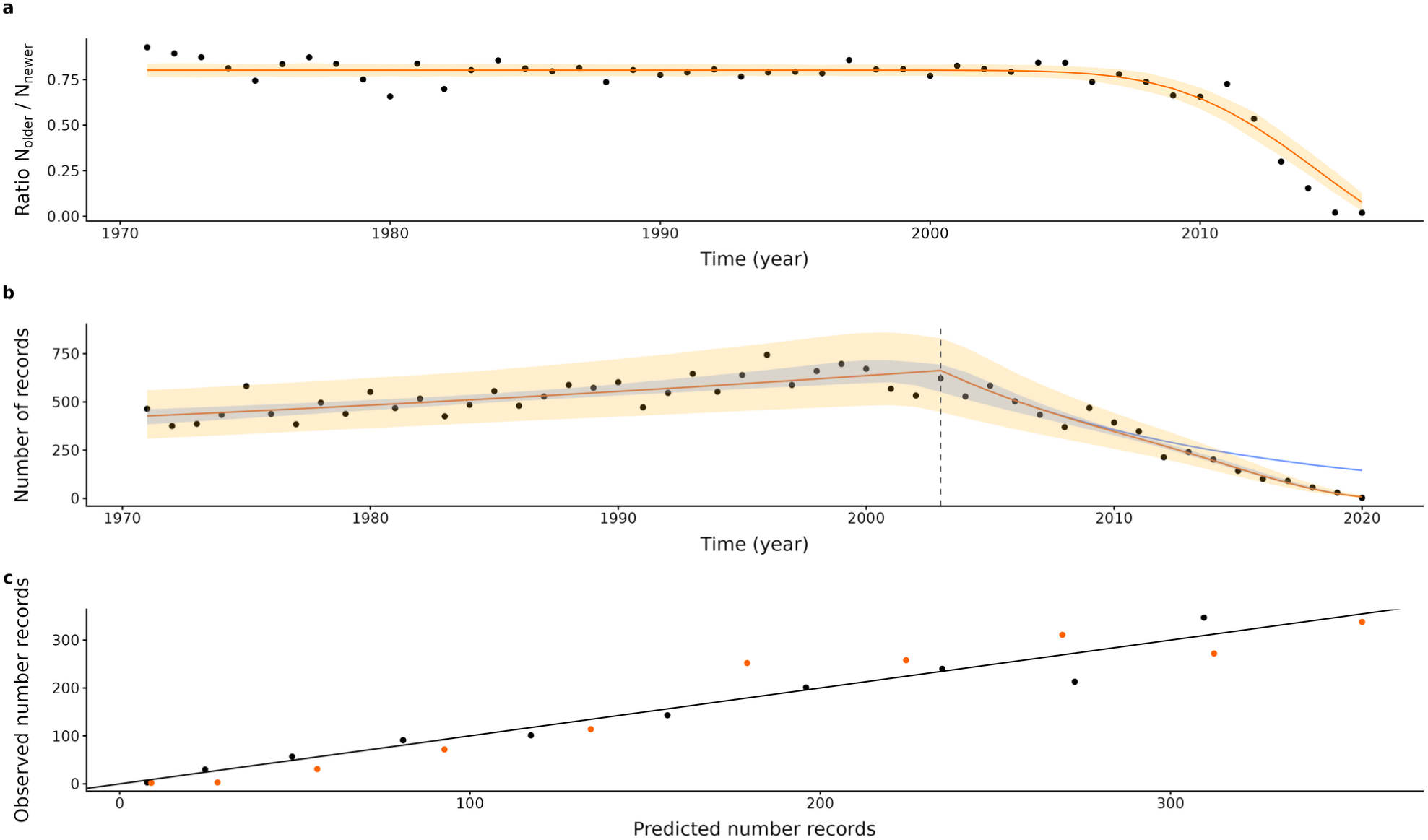
Fitted empirical patterns. a) Ratios of the 2017 and 2021 First Records databases (N_older_/N_newer_), comparing observed ratios (dots) versus the predictions from the *reporting ratio model* (line). b) Annual observed record counts (dots) versus predictions from the *record count model* (orange line), fitted on the 2021 database. Blue line indicates the estimated numbers of invasions, after accounting for reporting lags. Vertical line is the estimated change-point year. In both panels, credible intervals were estimated using the posterior distributions by refitting the *reporting ratio* and *record count models* with a Bayesian approach. Shaded area includes the upper and lower 95% Credible Intervals and additional stochastic variation from the observation model (using the 95% bounds from Binomial and Negative-Binomial, respectively). c) The predicted versus observed record counts during the 9-year estimated reporting lag. Black dots are from the 2021 database (r^2^=0.94), which was used to fit the model. Orange dots are from the 2017 database, estimated without refitting the model (r^2^=0.92), illustrating consistency in the reporting lag between 2021 and 2017, after accounting for publication date and differences in relative completeness.

Next, we used our *record count model* to examine invasion rates, using the 2021 database. Our results suggest that there was a change from increasing invasion rates to a decline in recent years, even after controlling for reporting lags (Fig 4b). Our best fitting model estimated that the numbers of invasions had been increasing at a rate of ∼1.5%/year, reaching an estimated maximum of 663 invasions in 2003, before subsequently declining to an estimated 146 invasions in 2020. Comparatively, there were only 3 empirically recorded invasions in 2020 (Fig 4b). We note that the low number of empirical records could reflect not only the strict definition of reporting lag (time between discovery and reporting), but also other temporal factors affecting inclusion in the database (e.g., change from classification as a casual to established invasion). Regardless of the mechanism, the important consideration is the effect of these factors on the temporal patterns of first records in the database, and that corrections are needed. Thus, the *record count model* indicated that the underlying invasion rate has substantially declined, but estimated that there was an order of magnitude higher numbers of invasions/year compared to the empirical records in 2020.

Models yielded significant improvements. For the *reporting ratio model*, we found a large improvement (ΔAIC = 1350.7) compared to the null including only differences in completeness between databases (i.e., no temporal reporting lag). For the *record count model*, we found ΔAIC = 96.6 against a constant invasion rate (no change point), suggesting a significant decline in invasion rate, beyond the decline due to the temporal reporting lag (Table 1). Further, the *record count model* substantially improved with the inclusion of reporting lag, compared to a null model differing only in completeness between databases (ΔAIC = 53.2, Table 1). Finally, the *record count model*, which was fit to 2021 database, worked comparably well when applied to the 2017 database (Fig 4c), despite markedly different numbers of invasion records per year. Thus, results from our global analyses suggest that the model operated as expected, and that including reporting lags and change-points were important in estimating the patterns of invasion records.

**Table 1:**
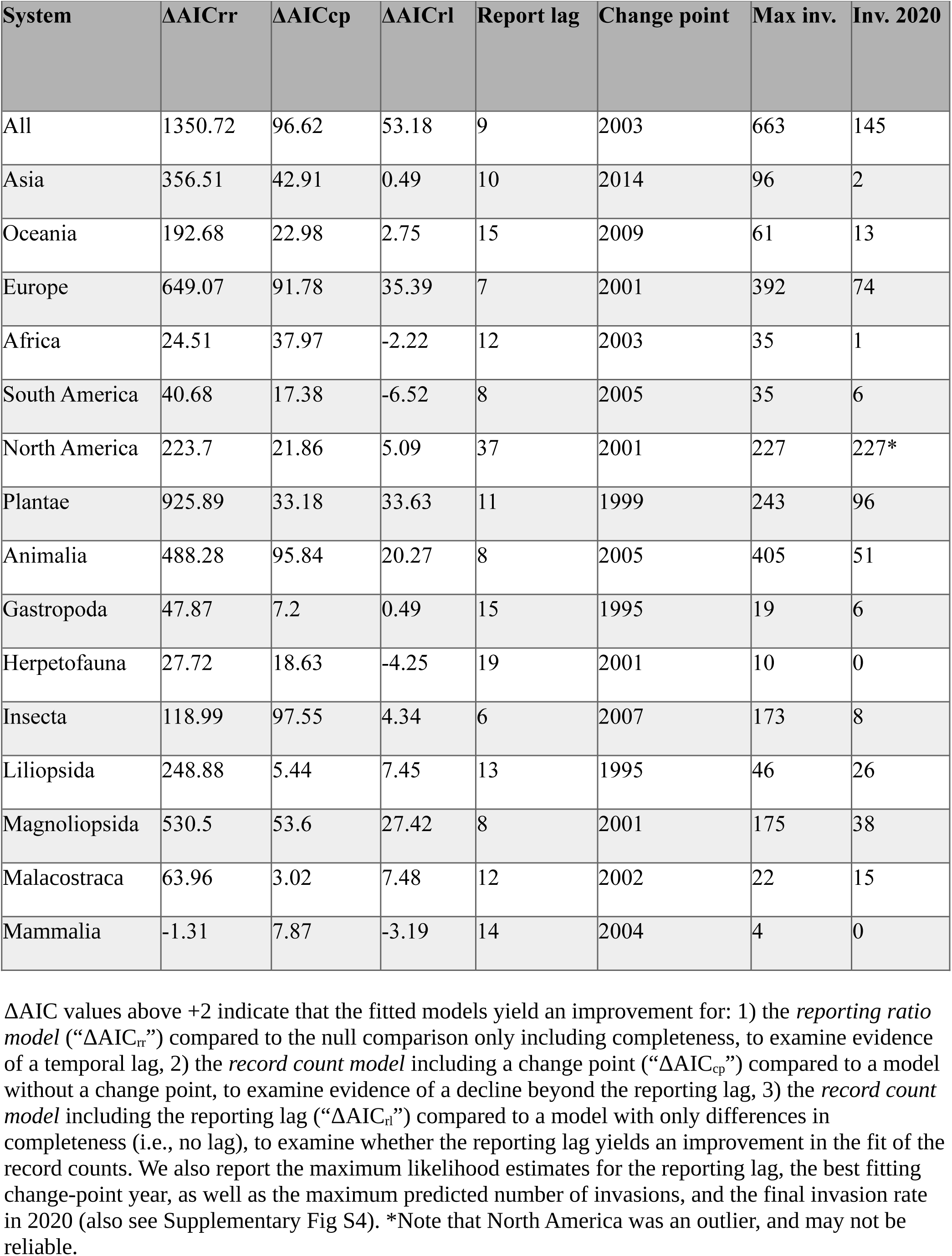
Regional and taxonomic sub-analyses. ΔAIC values above +2 indicate that the fitted models yield an improvement for: 1) the *reporting ratio model* (“ΔAIC_rr_”) compared to the null comparison only including completeness, to examine evidence of a temporal lag, 2) the *record count model* including a change point (“ΔAIC_cp_”) compared to a model without a change point, to examine evidence of a decline beyond the reporting lag, 3) the *record count model* including the reporting lag (“ΔAIC_rl_”) compared to a model with only differences in completeness (i.e., no lag), to examine whether the reporting lag yields an improvement in the fit of the record counts. We also report the maximum likelihood estimates for the reporting lag, the best fitting change-point year, as well as the maximum predicted number of invasions, and the final invasion rate in 2020 (also see Supplementary Fig S4). *Note that North America was an outlier, and may not be reliable.

| System | ΔAICrr | ΔAICcp | ΔAICrl | Report lag | Change point | Max inv. | Inv. 2020 |
|---|---|---|---|---|---|---|---|
| All | 1350.72 | 96.62 | 53.18 | 9 | 2003 | 663 | 145 |
| Asia | 356.51 | 42.91 | 0.49 | 10 | 2014 | 96 | 2 |
| Oceania | 192.68 | 22.98 | 2.75 | 15 | 2009 | 61 | 13 |
| Europe | 649.07 | 91.78 | 35.39 | 7 | 2001 | 392 | 74 |
| Africa | 24.51 | 37.97 | -2.22 | 12 | 2003 | 35 | 1 |
| South America | 40.68 | 17.38 | -6.52 | 8 | 2005 | 35 | 6 |
| North America | 223.7 | 21.86 | 5.09 | 37 | 2001 | 227 | 227* |
| Plantae | 925.89 | 33.18 | 33.63 | 11 | 1999 | 243 | 96 |
| Animalia | 488.28 | 95.84 | 20.27 | 8 | 2005 | 405 | 51 |
| Gastropoda | 47.87 | 7.2 | 0.49 | 15 | 1995 | 19 | 6 |
| Herpetofauna | 27.72 | 18.63 | -4.25 | 19 | 2001 | 10 | 0 |
| Insecta | 118.99 | 97.55 | 4.34 | 6 | 2007 | 173 | 8 |
| Liliopsida | 248.88 | 5.44 | 7.45 | 13 | 1995 | 46 | 26 |
| Magnoliopsida | 530.5 | 53.6 | 27.42 | 8 | 2001 | 175 | 38 |
| Malacostraca | 63.96 | 3.02 | 7.48 | 12 | 2002 | 22 | 15 |
| Mammalia | -1.31 | 7.87 | -3.19 | 14 | 2004 | 4 | 0 |

### Regional and taxonomic analyses

Next, we separated the database into six continental regions and considered nine taxonomic groups (animal and plant kingdoms, classes with more than 500 records since 1971, as well as vertebrate classes). We excluded Bacteria, Fungi and Protozoa, which accounted for less than 2% of all the records). Additionally, we excluded analyses of Aves - reporting lags were estimated to be very long (centuries), which could arise as an artifact of higher reporting probability in the latter database (see section “*Theoretical* e*ffects of changes in effort*”), showing no evidence of an asymptotic difference between databases. Finally, we observed several outliers (one for Oceania and Gastropoda, and three for Mammalia) (see Supplementary Fig S4). These spikes in records had the same patterns as theoretically expected with a short-term increase in effort (compare with Supplementary Fig S3), and thus were removed prior to analyses.

The number of recorded entries in the First Records database since 1970 (22K total) was dominated regionally by Europe (with >11K), and thus, the global analyses above could have largely reflected patterns in Europe. However, we found that most sub-analyses showed broadly similar patterns (Table 1) - the estimated reporting lags generally centered around a decade, with Oceania and Gastropods showing a 15-year lag, and minimums for Europe (7 years) and Insecta (6 years). However, there were two exceptions, for North America (37 years) and Herpetofauna (reptiles and amphibians - 19 years) (Table 1). Further, all sub-analyses found improvements for the *record count model* (using ΔAIC compared to a null model that did not include a change-point), and all but mammals showed improvements in the reporting ratio model (compared to null models that only accounted for differences in completeness) (Table 1). Further, the best-fitting models estimated that all regions and taxa had increasing rates of invasion until a change point, and almost all showed subsequent declines, with North America as the sole exception (Supplementary Figure S4). Thus, the most parsimonious interpretation is that there was a true declining rate of invasion in most regions and taxa.

## Discussion

Our results suggest that reporting lags are important, that there remains substantial uncertainty for some regions and taxonomic groups, but that real declines in the invasion rate have occurred over the last two decades. These findings could dramatically alter previous projections of rates of biological invasions. Recent analyses projected increasing trends but omitted the last two decades of observations (Seebens et al. 2025, IPBES 2023). While this was logical given the potential effect of reporting lags, and our analyses agree that rates were increasing up to the early 2000s, our current work allows these analyses to be updated to include and interpret recent declining trends.

Additionally, our findings suggest that many previous analyses projecting invasions rates should be revisited (including some of our own work; e.g., Sardain et al. 2019, Leung et al. 2014). Previous projections were parameterized based on historical invasion patterns, often preceding the last two decades. These invasion patterns were then related to different direct and indirect drivers, such as global shipping (e.g., Sardain et al. 2019), trade (e.g., Brockerhoff et al. 2014), and GDP (e.g., Leung et al. 2014). Given the putatively positive relationship between these drivers and propagule and colonization pressures (Lockwood et al. 2009), and the primacy of these processes for invasions, these previous projections have all been rational. However, while these drivers have all broadly increased over the last two decades, our current results suggest the number of invasions has declined. This suggests model predictions need to be reevaluated, that the decline in invasions could be even more acute based on a per unit of introduction pressure, and that other broad factors are at play.

One explanation for the decline in invasion rate could be the expansion of policies aimed at reducing or mitigating invasions over recent decades (Canelles et al. 2025), the consequences of which could be to reduce the effective propagule pressure despite increases in anthropogenic drivers. If this were the case, the decline in invasion rate would provide support for recent efforts to reduce invasions (e.g., EU Regulation 1143/2014, European Commission 2014). Alternatively, an additional mechanism could include depletion of invasion source pools that can be plausibly associated with particular pathways and vectors, particularly as inter-regional spread saturates (Seebens et al. 2021). Although, as climatic and geopolitical changes alter trade pathways, invasion probabilities could concomitantly shift by connecting novel source pools.

More theoretically, other work has largely focused on discovery (detection) lags (Solow & Costello 2004, Robeck et al. 2024). In contrast, our work pointed to reporting lags as a critical factor for interpreting recent trends based on invasion counts. This occurs because the number of invasions discovered per year may eventually equilibrate with the number of undiscovered invasions, allowing the underlying rate of invasion to be estimated. However, we note that discovery lags remain important in other contexts - for early invasions, or estimating the timing of individual species’ invasions.

While recognizing that the First Records database is widely used to estimate invasion rates, and is arguably the best dataset that we have to do so, our analyses show that real declines in invasion rate have been happening, while highlighting interpretation challenges for these compilation databases. For instance, comparisons of all databases did not show any signature that the decline in invasions was due to lags, as in such a case, we would have expected a shift and increase in the peak rate for more recent databases (instead the peak stayed virtually constant). However, the most recent 2023 and 2025 databases also showed few additional records after 2020, with zero reported records after 2021 and 2022, respectively, and noticeably less effect of reporting lag than the 2017 versus 2021 database versions. This could reflect lower reporting effort in the recent databases, for instance as a consequence of the COVID19 pandemic as one possible cause, which would result in the trend being more strongly attributed to true declines in invasion rates. Thus, our analyses using 2017 and 2021 is the more conservative estimate of decline rates, if one is to infer trends using the First Records databases, and it still shows substantial decline. Moreover, other metrics likewise provide confirmatory evidence for using the 2017 versus 2021 comparisons - namely, predictions built using 2021 were strongly predictive of 2017 patterns, despite a strong shift in numbers for a given year during the lag phases, and inclusion of the lag from the *reporting ratio model* yielded substantial improvements in the observed record counts.

Additionally, the best-fitting trends were qualitatively similar between most sub-analyses, although our confidence varied for different regions and taxa. While fit to the *record count model* improved by adding a reporting lag for most analyses, several groups (i.e., regions: Africa, Asia, South America; taxa: Gastropoda, Herpetofauna, Mammalia) did not show improved AIC values. This reflected relatively high variation in the datasets of those regions and taxa, and thus less confidence regarding the estimated rate of decline (see Supplementary Fig S2). Additionally, North America was an outlier both in terms of a long estimated reporting lag (the difference between database versions continued to increase across decades, in contrast to all other regions which generally reached an asymptote quickly), and the consequent estimated increase in invasion rates (all other regions predicted declines). Other evidence of problems with the North American signal include the fact that only <10% of records were from North America compared to 50% from Europe, whereas other evidence suggests the numbers of invasions should be similar between the two continents (Briski et al. 2024). Thus, we view some of the regional and taxonomic relations cautiously.

It is prudent to highlight that our analyses only focused on trends for novel invasions in regions of substantial size (countries, islands). Importantly, intra-regional spread and ecological or socio-economic impacts will continue to expand and accumulate (e.g., Liu et al. 2023, Liu et al. 2025), even considering only alien species that have already arrived and established (of course, novel invasions also continue to occur, albeit at a slower rate). Thus, management options (e.g., eradication, ‘slow the spread’, and risk analyses, Lodge et al. 2006), and empirical evaluation of policy effects on global, regional and taxonomic trends remain relevant (e.g., is the rate of intra-regional spread slowing? Can we empirically observe differences in spread rates or their impacts, caused by different policies?). In this regard, databases such as GBIF (for intra-regional spread, GBIF.org 2025) and InvaCost (for economic impacts, Diagne et al. 2021) already exist and could be useful. However, as seen in our analyses of invasion rates, interpretation of trends using such compilation databases can be challenging and nuanced. Regardless, continued management, evaluation of trends and potential identification of opportunities for improvements at all invasion stages remain pertinent.

Irrespective of other invasion stages and possible additional analyses, the potential ramifications of our current findings are broad and timely. The Global Biodiversity Framework (CBD 2022) aims to “bend the curve” and reduce the rate of invasions by 50% by 2030. It may actually be that the policies that have been put in place, including ballast water regulation (IMO 2004), International Sanitary and Phytosanitary Measures (e.g., ISPM15, IPPC Secretariat 2009), the European Union IAS Regulation (European Commission 2014), as well as education and awareness campaigns (Haley et al. 2023), have brought us closer to those targets. Importantly, this would not suggest that the invasions are no longer an issue and invasion policies no longer relevant - discontinuing management efforts would likely result in a reversal back to historical invasion rates and a greater escalation of impacts. Regardless, our findings provide positive news, and suggest that the future for stemming the tide of biological invasions could look more promising than previously expected.

## Acknowledgements

We thank Hanno Seebens for input on this manuscript, as well as members of the Leung Lab for discussions. This work was funded by a Natural Sciences and Engineering Research Council Discovery grant and Fonds de recherche du Québec Société et culture (https://doi.org/10.69777/337747) grant to BL.

## Notes

### Competing Interest Statement

The authors have declared no competing interest.

