## Supplementary Information for "Recent trends show a 20 year decline in global biological invasion rates"

**Table S1: Glossary**

| Term | Description |
|---|---|
| Alien species | A species that is introduced from one region to another region, where it does not naturally occur, typically via human activities. The designation as alien species is irrespective of any impacts. |
| Biological invasion | The process by which a species is introduced into another region, where it does not naturally occur, typically via human activities. The species can potentially establish, spread and cause impacts in the new region, as part of the overall invasion process. However, in the context of this study, our focus on biological invasions is restricted to alien species which have established. |
| Change point | In the context of this study, a time point (year) in which the rate of invasion changes (equation 2). This can include rate changes that become more positive or negative, as well as changing directions (e.g., from an increasing to a decreasing rate). |
| Database completeness | The fraction of reported invasions that are included in a database (denoted by $M_T$, equation 7, in the main text). To be included, an invasion must first be discovered and reported. |
| Discovery lag | The time period between an invasion event and discovery. |
| Discovery probability | The probability that an invasion will be discovered per time interval (year), after an invasion event. Discovery probability can be constant or can increase as an invasion progresses, up to a maximum probability, which are both included in equation 3. Discovery probability less than one can result in a discovery lag. |
| First Record | The first recorded instance of a species in a given region, forming the data in the First Records Database (Seebens *et al.* 2017). To become a First Record, an invasion must be discovered, reported, and subsequently entered into the First Records database. Not all reported invasions will be entered into the database (database completeness), which affects the record count (equation 7). |
| Invasion rate | The rate that new species are introduced into new regions (equations 1 & 2). In the model, we consider an initial invasion rate (equation 1), as well as a change in invasion rate, after a change point (equation 2). In the context of the First Records database, this includes only invasions which become established. |
| Record count model | The model estimating parameters for the invasion process, by comparing observed versus expected number of records over time, after adjusting for reporting lag (output from the *reporting ratio* model described below), using maximum likelihood (equation 10). |
| Reporting lag | The time period between discovery of an invasion and reporting. Note, not all |

| | |
|---|---|
| | reported invasions will be entered into a database (i.e., databases are incomplete). To provide a single number, reporting lag is calculated as the number of years before >95% of invasions have been reported, reflecting the *relative completeness* within a database. This distinction is made because absolute levels of database completeness are unknown. |
| Reporting probability | The probability that an invasion will be reported per time interval (year), after discovery. Reporting probability can be constant or can increase as an invasion progresses, up to a maximum probability (equation 5). Reporting probability less than one can result in a reporting lag. |
| Reporting ratio model | The model estimating parameters for the reporting lag and relative database completeness. This is done by comparing the record counts from two database versions using maximum likelihood (equation 8 & 9). We note that the model captures temporal effects of reporting lag, but also other temporal factors affecting inclusion in the database (e.g., change in classification from casual to established invasions). Regardless of the mechanism, temporal relationships need to be estimated to interpret relative trends *within* a database. The model does this, after controlling for general, non-temporal, differences in database completeness, required because the model compares *between* databases. |

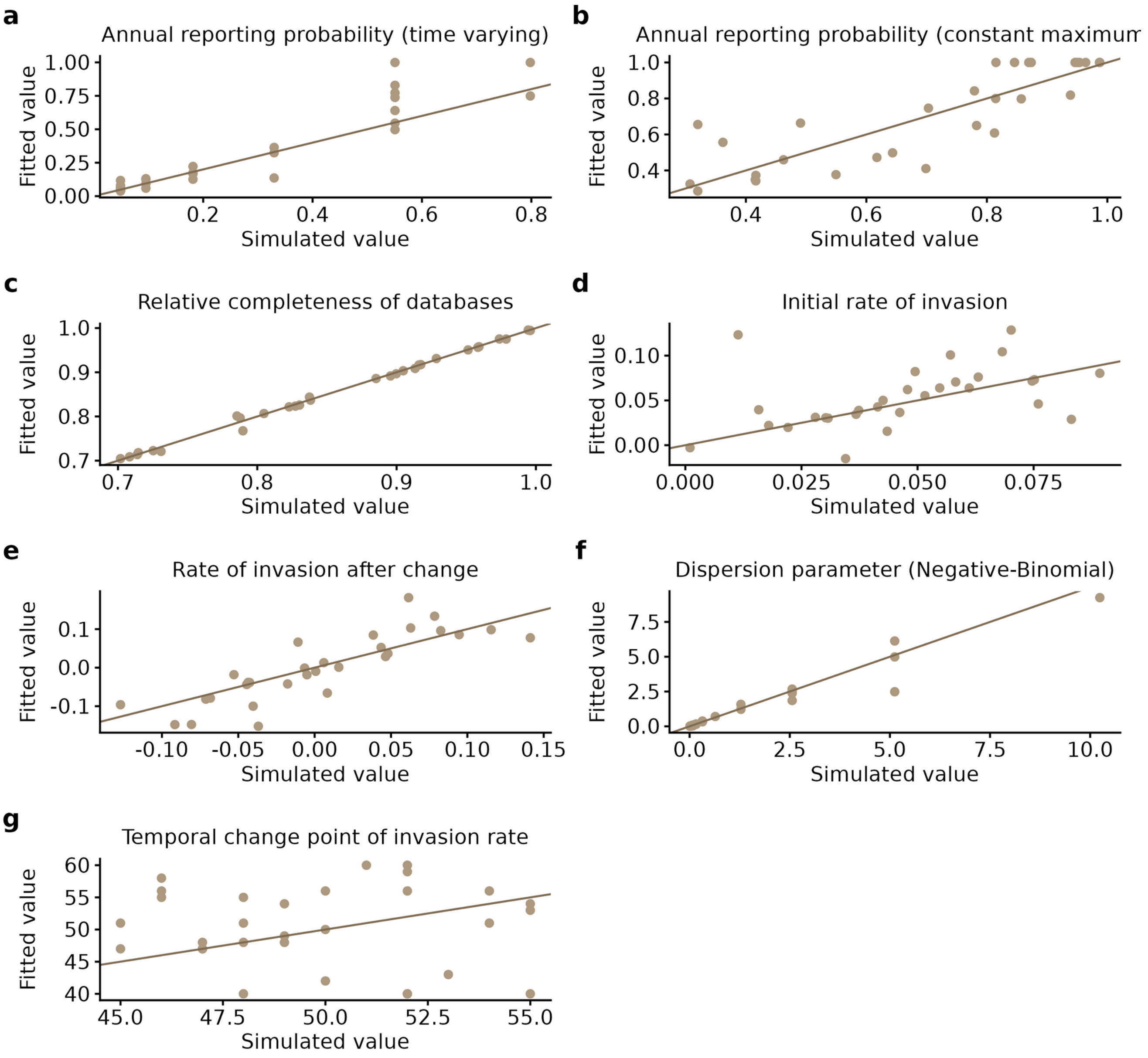


**Figure S1.*Theoretical analyses using simulations.*** Panels a-c: evaluation of *reporting ratio model*, comparing simulated (equation 5) versus fitted probabilities via maximum likelihood (equation 9), for a time-varying (panel a) and maximum (panel b) annual reporting probability (corresponding to parameters $\alpha$ and $\beta$, respectively). The time-varying component of probability is presented as the first year (t=1) after discovery ($1-e^{-\alpha t}$) (panel B). As $1-e^{-\alpha t}$ approaches one, we approach a constant maximum annual probability of reporting ($\beta$, Panel b). Panel c shows relative completeness (parameter $M_R$, equation 8. Panels d-g: evaluation of *record count model*, comparing simulated (equation 1-2) versus fitted parameters using maximum likelihood (equations 10), after controlling for reporting lags. The approach was able to estimate the changes in invasion rates (panels d, e), as well as the sampling variation (the dispersion parameter from a Negative-Binomial distribution, panel f), but not the change point year (panel g). The model (equation 10) was modified to exclude discovery lags, which could not be reliably estimated (Supplementary Fig S2) but was not needed to estimate invasion rates. One-to-one lines indicate a perfect estimation between fitted and simulated values (solid black lines).

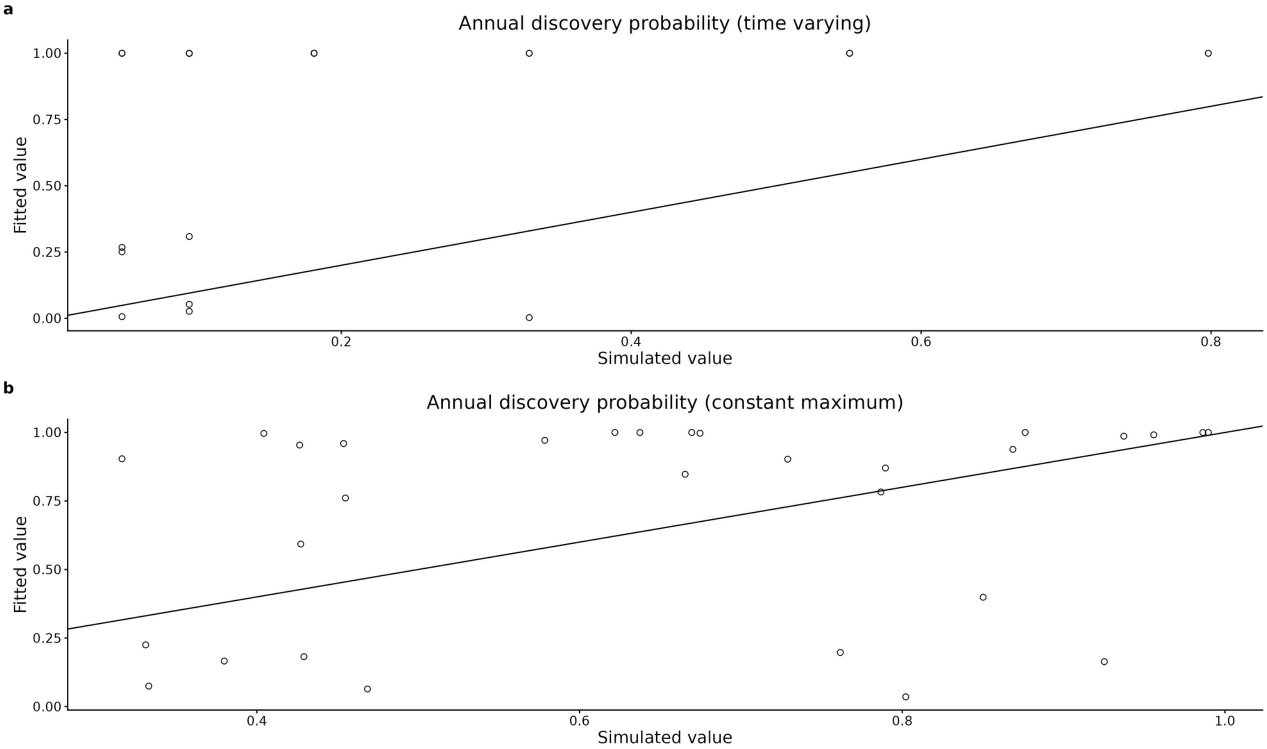


**Figure S2. Theoretical analyses of discovery probability**. Simultaneous fit of discovery probabilities with invasion rates was not able to correctly estimate either the simulated time varying component (Panel a) nor the maximum discovery probability (Panel b). This could occur because discovery rates would equilibrate with invasion rates over time, irrespective of the discovery probability, such that there would be insufficient information to separate out each independent effect. However, for this same reason, one could ignore discovery probability and still accurately estimate invasion rates, which was the final procedure conducted in the main text.

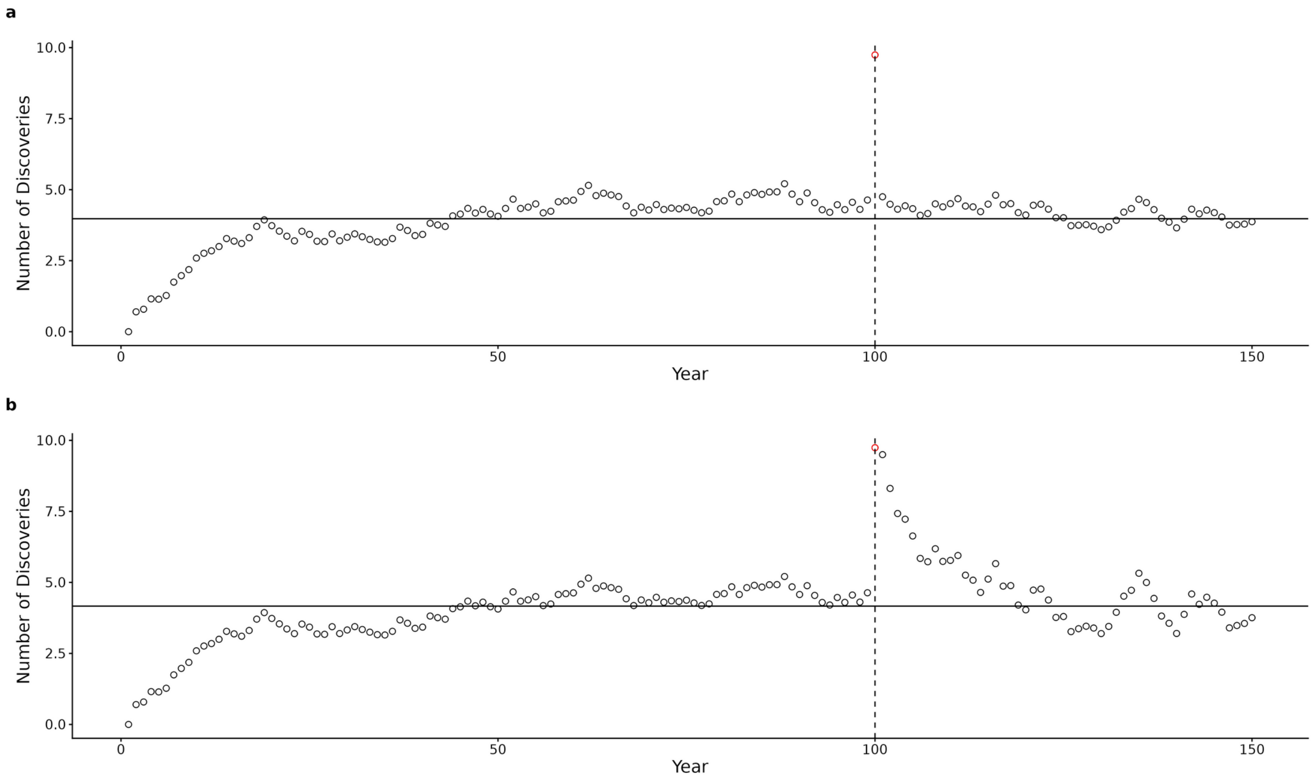


**Figure S3.** ***Theoretical Effect of changing effort.*** The model assumes that the discovery and reporting probabilities remain relatively consistent across the time series. This may be violated for instance if research effort strongly changes. However, changes in effort can result in characteristic signals. A single one year pulse of research effort would result in a spike in discoveries (red dot) followed by an immediate decline back to equilibrium (Panel a). A persistent increase in research effort would also result in pulse, but would still result in a slower decline back to equilibrium, where discovery rate equals invasion rate (see main text for explanation). This characteristic signal – a spike in discoveries - can help identify when patterns are due to changes in effort rather than true changes in invasion rate.

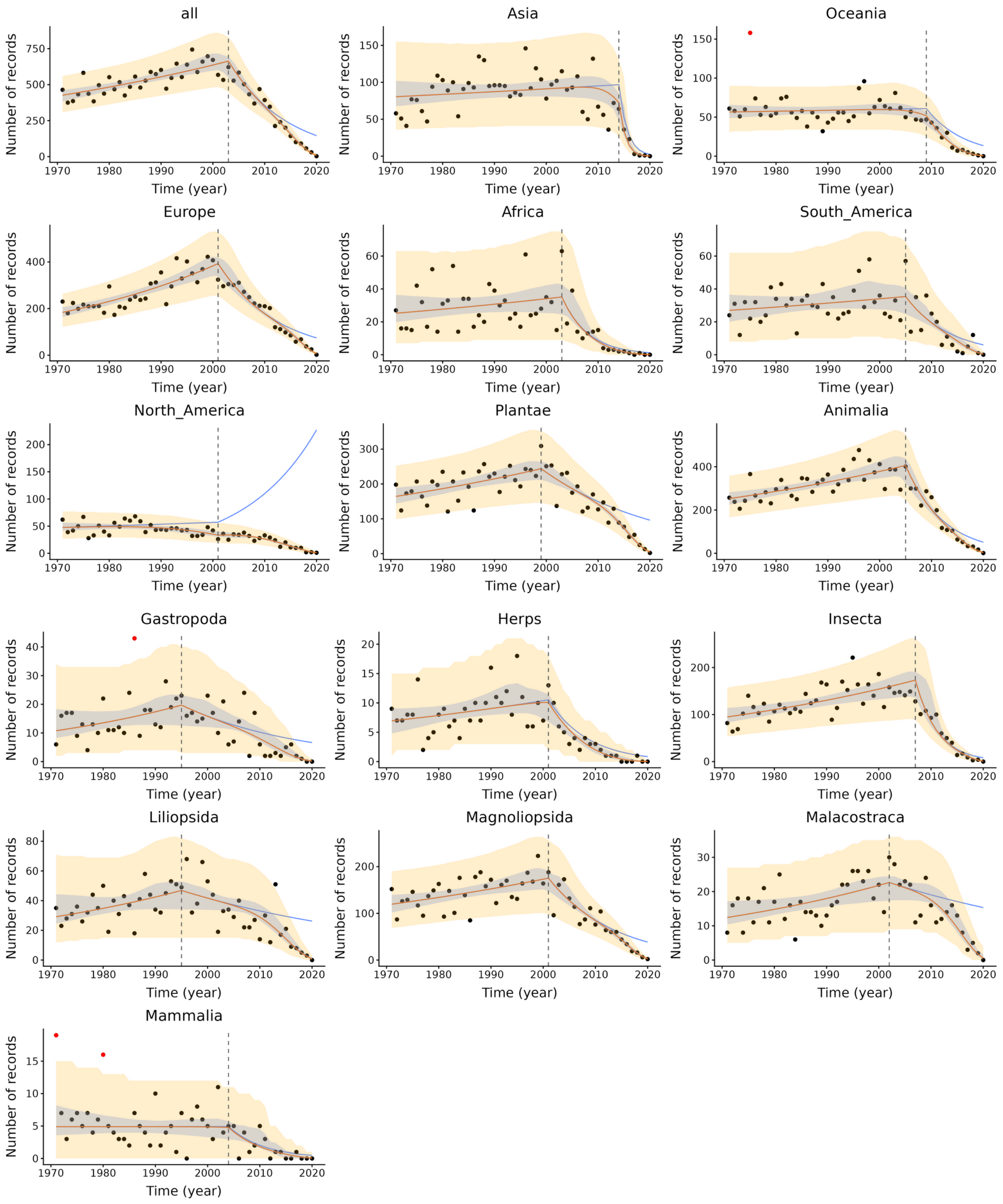


**Figure S4. Regional and taxonomic patterns**. Annual observed record counts (dots) versus predictions from the *record count model* (orange line), fitted on the 2021 database. Shaded area includes the upper and lower 95% Credible Intervals and additional stochastic variation using the 95% bounds from a Negative-Binomial observation model. Blue line indicates the estimated numbers of invasions, after accounting for reporting lags. Vertical line is the estimated change-point year. Red dots are outliers, similar to characteristic spikes expected due to a pulse of research effort (see Supplementary Figure S3a) rather than higher invasion rate, and thus were removed.

**Supplementary discussion of regional and taxonomic sub-analyses**

The best-fitting trends were qualitatively similar between most sub-analyses, however, our confidence varied for different regions and taxa. We looked for indicators of potential problems for interpretations, to identify where additional caution was warranted. While the regional patterns were largely as expected, with Global South countries showing more variability in first records patterns, we had expected vertebrates to show clearer signals than other groups, given conspicuousness and broad societal interest. Instead, vertebrates were generally less certain across a number of metrics, from a non-significant *reporting ratio model* (for mammals), long reporting lags (for Herpetofauna and Aves) and greater variation in data points. Potentially, this could in part be due to relatively low numbers of records, for these regions and taxa, which should thus be viewed with caution. Additionally, we were uncertain about the North American trend, where the model estimated a 37-year reporting lag. Given the economic growth and globalization, especially of the United States, it may make sense that North America stands as an outlier. Conversely, such a long reporting lag is surprising given that North America is generally considered well-funded and data rich. An alternative possibility is that there was an *increase* in reporting probability between the 2017 and 2021 databases inflated the apparent reporting lag, causing an artificially high invasion trend (logic explained in *"theoretical effects of changes in effort"* in main text). Mechanistically, the long lag occurred because the difference in records between databases did not clearly reach an asymptote, in contrast to other regions. A further indication of some reporting oddities is in the relative number of records between databases: other compilations of invasion records suggest that Europe and North America both have comparably large fractions of known invasions (Briski et al. 2024). In contrast, in the First Records database, only 1.8K (<10%) of the records were from North America, while 50% was from Europe. Thus, while the long reporting lag and rate of increase in invasions are plausible, the estimates for North America should be viewed with caution, and require further exploration.
